# A map of direct TF-DNA interactions in the human genome

**DOI:** 10.1101/394205

**Authors:** Marius Gheorghe, Geir Kjetil Sandve, Aziz Khan, Jeanne Chèneby, Benoit Ballester, Anthony Mathelier

## Abstract

Chromatin immunoprecipitation followed by sequencing (ChIP-seq) is the most popular assay to identify genomic regions, called ChIP-seq peaks, that are bound in vivo by transcription factors (TFs). These regions are derived from direct TF-DNA interactions, indirect binding of the TF to the DNA (through a co-binding partner), nonspecific binding to the DNA, and noise/bias/artifacts. Delineating the bona fide direct TF-DNA interactions within the ChIP-seq peaks remains challenging. We developed a dedicated software, ChIP-eat, that combines computational TF binding models and ChIP-seq peaks to automatically predict direct TF-DNA interactions. Our work culminated with predicted interactions covering >4% of the human genome, obtained by uniformly processing 1,983 ChIP-seq peak data sets from the ReMap database for 232 unique TFs. The predictions were a posteriori assessed using protein binding microarray and ChIP-exo data, and were predominantly found in high quality ChIP-seq peaks. The set of predicted direct TF-DNA interactions suggested that high-occupancy target regions are likely not derived from direct binding of the TFs to the DNA. Our predictions derived co-binding TFs supported by protein-protein interaction data and defined *cis*-regulatory modules enriched for disease- and trait-associated SNPs. Finally, we provide this collection of direct TF-DNA interactions and *cis*-regulatory modules in the human genome through the UniBind web-interface (http://unibind.uio.no).

## INTRODUCTION

The transcription of DNA into RNA is mainly regulated through a complex interplay between proteins and the chromatin at *cis-*regulatory regions such as promoters and enhancers. Transcription factors (TFs) are key proteins specifically binding short DNA sequences, known as TF binding sites (TFBSs), to ensure transcription at appropriate rates in the correct cell types (1). Therefore, genome-wide identification of TFBSs is a critical step to decipher transcriptional regulation, and how this process is altered in diseases (2).

Classically, genome-wide *in vivo* TF binding regions are identified through the chromatin immunoprecipitation followed by sequencing (ChIP-seq) assay (3). The genomic regions obtained with ChIP-seq, the so-called ChIP-seq peaks, are usually a few hundred base pairs (bp)-long and should encompass the TFBSs (∼10 bp-long), where direct TF-DNA interactions occur. However, ChIP-seq peaks derive from either direct TF-DNA interactions, protein-protein interactions with other regulators such as co-factors, or unspecific binding. Moreover, ChIP-seq experiments are prone to artifacts and delineating bona fide TF-bound regions is still an ongoing challenge (4–7).

As TFs specifically recognize DNA sequence motifs, computational tools have been instrumental in the prediction and characterization of direct TF-DNA interactions (8). TFBSs are commonly modelled with position weight matrices (PWMs), which represent the probability of each nucleotide to be present at each position within bona fide TFBSs (8). While PWMs work well (9), more sophisticated approaches have recently been designed to model complex features of TF-DNA interactions captured by next-generation sequencing data (10). However, the best performing model varies for different TFs or TF families (9, 11, 12).

While multiple resources collecting TF binding regions derived from ChIP-seq exist (13–16), a limited number store genome-wide identification of TFBSs (14, 17, 18). The TFBS Conserved Track of the UCSC Genome Browser combined phylogenetic sequence conservation and PWMs to identify TFBSs (19) while the MANTA resource (20) integrated ChIP-seq peaks from ReMap (13) with PWMs from JASPAR (21) for TFBS predictions. A strong limitation of these approaches is that they use the same pre-defined score thresholds for all PWMs and all data sets. The ORegAnno database provides TFBSs obtained through literature curation (18, 22), but the number of TFBSs available for human is limited to only ∼8,000.

A previous study showed that ChIP-seq data sets fall within one of three categories: (i) data sets enriched for the TF canonical binding motif close to the ChIP-seq peak summit (where the highest number of ChIP-seq reads map), (ii) data sets lacking enrichment for the canonical binding motif close to the peak summit, and (iii) data sets having a combination of peaks with and without the TF canonical binding motif proximal to the peak-summit (23). Most ChIP-seq data sets were observed in category (iii). As direct TF-DNA interactions are expected to be enriched at ChIP-seq peak summits (23–28), Worsley Hunt *et al.* developed a heuristic approach specifically based on PWMs to automatically identify, in each ChIP-seq data set, this enrichment zone. The method determines the thresholds on the PWM scores and distances to the peak summits delimiting the enrichment zone that contains direct TF-DNA interactions. However, this method does not work with the more recent TFBS computational models (12, 29, 30).

In this study, we mapped direct TF-DNA interactions in the human genome in a refined manner by capitalizing on uniformly processed TF ChIP-seq data sets and computational tools modelling TFBSs. We provide (i) a new software to predict direct TF-DNA interactions within ChIP-seq peaks along with (ii) genome-wide predictions of such interactions in the human genome. Using an entropy-based algorithm, we have developed ChIP-eat, a tool that automatically identifies direct TF-DNA interactions using both ChIP-seq peaks and any computational model for TFBSs. We applied ChIP-eat to 1,983 human ChIP-seq peak data sets from the ReMap database (13), accounting for 232 distinct TFs. The set of predicted direct TF-DNA interactions derived from PWMs covers >4% of the human genome. To make this resource available to the community, we have created UniBind (http://unibind.uio.no/), a web-interface providing public access to the predictions. We validated *a posteriori* these TFBS predictions using protein binding microarray (31) and ChIP-exo (32) data, and multiple ChIP-seq peak-callers. We used these TFBSs to (i) confirm that hotspots of ChIP-seq peaks (also known as high occupancy target regions (33)) are likely not derived from direct TF-DNA interactions, (ii) predict co-binding TFs, and (iii) define *cis*-regulatory modules, which are enriched for disease- and trait-associated SNPs.

## MATERIALS AND METHODS

### ChIP-seq data

The ChIP-seq data sets considered were retrieved, processed, and classified as part of the last update (2018) of the ReMap database (13) (Supplementary Figure S1).

### TF binding profiles

For 1,983 ChIP-seq data sets used in the last ReMap update, we were able to manually assign TF binding profiles corresponding to the ChIP’ed TFs as position frequency matrices (PFMs) from the JASPAR (2018) database (21).

### Training data sets

To train the TFBS computational models (see below), we considered 101 bp sequences centered around the peak summits as positive training sets. When required for training, negative training sets were obtained by shuffling the positive sequences using the *g* subcommand of the BiasAway (version 0.96) tool to match the %GC composition (23).

### TFBS computational models

#### Position weight matrices

JASPAR PFMs were converted to PWMs as previously described in (34). For each ChIP-seq data set, PWMs were optimized using DiMO (version 1.6; default parameters with a maximum of 150 optimization steps) using the corresponding training sets (35).

#### Binding energy models

JASPAR PFMs were converted to binding energy models (BEMs; (30)) using the implementation from the MARS Tools (https://github.com/kipkurui/MARSTools; (36)). We modified the implementation to return a BEM score corresponding to 1 - (original score) to consider the best site of the DNA sequence as the one with the highest BEM score (instead of the lowest one).

#### Transcription factor flexible models

First-order transcription factor flexible models (TFFMs) (version 2.0) were initialized with the DiMO-optimized PFMs and trained with default parameters (https://github.com/wassermanlab/TFFM; (29)) on the positive training sets.

#### DNAshapedTFBS models

The DNA shape-based models were trained on the training sets using the DNAshapedTFBS tool (version 1.0; https://github.com/amathelier/DNAshapedTFBS/; (12)). We trained three types of DNAshapedTFBS models with the following features: (i) DiMO-optimized PWM + DNA shape, (ii) first-order TFFM + DNA shape, and (iii) 4-bits encoding + DNA shape following (12). We considered the first and second order DNA shape features helix twist, propeller twist, minor groove width, and roll with values extracted from GBShape (37).

### Landscape plots

Each TFBS computational model was applied to each ChIP-seq data set independently. Following the strategy described in (23), we considered 1,001 bp sequences centered around the peak summits, obtained using the bedtools (version 2.25) *slop* subcommand (38). The trained computational models were used to extract the best (maximal score) site per 1,001 bp ChIP-seq peak region. For each ChIP-seq data set, landscape plots were constructed from the corresponding sites following the TFBS_Visualization tool (23). These scatter plots were also converted into heat maps using the *kde2d* function from the MASS R package (39).

### Automated identification of the enrichment zone

To define the enrichment zone for each landscape plot, we automatically identified the thresholds for the TFBS computational model scores and distances to peak summits using the entropy-based algorithm from (40). The algorithm aims at identifying two classes of elements. Given a histogram, the algorithm selects the threshold that maximizes the within-class sum of the Shannon entropies for the elements in two classes (41). The two classes of elements identified are defined by the elements with values i) above and ii) below the threshold, respectively. This procedure optimally separates the input elements in two classes. Given a ChIP-seq data set, we applied the algorithm to the histograms of the TFBS computational model scores and distances to peak summits, independently. The maximum entropy implementation of the algorithm available in ImageJ (42) was used with default parameters.

The source code of the ChIP-eat software used to process ChIP-seq peak data sets to predict direct TF-DNA binding events is freely available at https://bitbucket.org/CBGR/chip-eat. Specifically, ChIP-eat trains a TFBS computational model and automatically defines the enrichment zone in the landscape plots to predict the underlying direct TF-DNA interactions. The identification of the enrichment zone has been applied to each TF ChIP-seq peak data set independently, allowing for the automatic detection of the thresholds that are specific to each data set with each TFBS computational model.

### Assessing the robustness of the enrichment zone identification

For each ChIP-seq data set, we sampled the set of peaks using the seqtk (version 1.0) (https://github.com/lh3/seqtk) *sample* subcommand. The sequences of the sampled peaks were shuffled using the *fasta-shuffle-letters* subcommand of the MEME suite (version 4.11.4) (43) and added to the original set of ChIP-seq peaks. The automatic thresholding was applied to this new set. We tested the addition of shuffled peaks representing 10%, 25%, and 50% of the original set peaks.

### Comparison with the heuristic approach to predict the enrichment zone

ChIP-eat was compared to the heuristic approach described in (23) and implemented in the TFBS_Visualization tool https://github.com/wassermanlab/TFBS_Visualization using the default parameters. The centrality of the TFBSs predicted in the enrichment zones predicted by ChIP-eat and TFBS_Visualization was assessed using centrality p-value computations as described in the CentriMo tool (25).

### TF-DNA binding affinity assessment with protein binding microarray data

Protein binding microarray (PBM) (44) data were retrieved from UniProbe (http://the_brain.bwh.harvard.edu/uniprobe/; (45)) for 40 TFs with available ChIP-seq data. For each ChIP-seq data set landscape plot, we extracted the DNA sequences at the sites within and outside of the predicted enrichment zone. The binding affinity of a TF to each site was computed as the median PBM intensity value of all the de Bruijn sequences containing the site sequence. The statistical difference between the distribution of PBM binding affinities from sites within and outside the enrichment zone was assessed using a two samples Mann-Whitney U test (46) implemented in the R package *stats*. A Bonferroni correction was applied to the computed p-values.

### ChIP-exo data

ChIP-eat was applied with DiMO-optimized PFMs to the ChIP-exo data sets from (47), which were lifted over to hg38 using the liftOver tool (17). As for ChIP-seq peaks, we considered 1,001 bp regions centered around the peak summits.

### ChIP-seq peaks from HOMER and BCP peak-callers

We successfully applied the HOMER (version 4.7.2) (48) and BCP (version 1.1) (49) peak-callers to 670 ENCODE ChIP-seq data sets (Supplementary Table S1). ChIP-eat was applied to the corresponding ChIP-seq peak regions with DiMO-optimized PFMs as described above. ChIP-seq peaks predicted to contain a direct TF-DNA interaction or not (using the enrichment zones) from the three peak-callers (MACS2 (50), HOMER, and BCP) were overlapped using the bedtools *intersect* subcommand. Hypergeometric tests were performed to assess the significance of the intersections using the R *phyper* function for every combination of two peak-callers with the following contingency matrix:

|  |  |
| --- | --- |
| number of overlapping peaks **with** TFBSs from two peak-callers - 1 | number of peaks **without** TFBSs from the two peak-callers |
| number of peaks **with** TFBSs from the two peak-callers | number of overlapping peaks from the two peak-callers |

### HOT/XOT regions

The high occupancy target (HOT) and extreme occupancy target (XOT) regions in all contexts were downloaded through the ENCODE data portal at http://encode-ftp.s3.amazonaws.com/modENCODE_VS_ENCODE/Regulation/Human/hotRegions/maphot_hs_selection_reg_cx_simP05_all.bed and http://encode-ftp.s3.amazonaws.com/modENCODE_VS_ENCODE/Regulation/Human/hotRegions/maphot_hs_selection_reg_cx_simP01_all.bed (51)). ChIP-seq peaks were overlapped with the HOT/XOT regions using the bedtools *intersect* subcommand. The enrichment for overlap was assessed with a hypergeometric test using the R *phyper* function with the following contingency matrix:

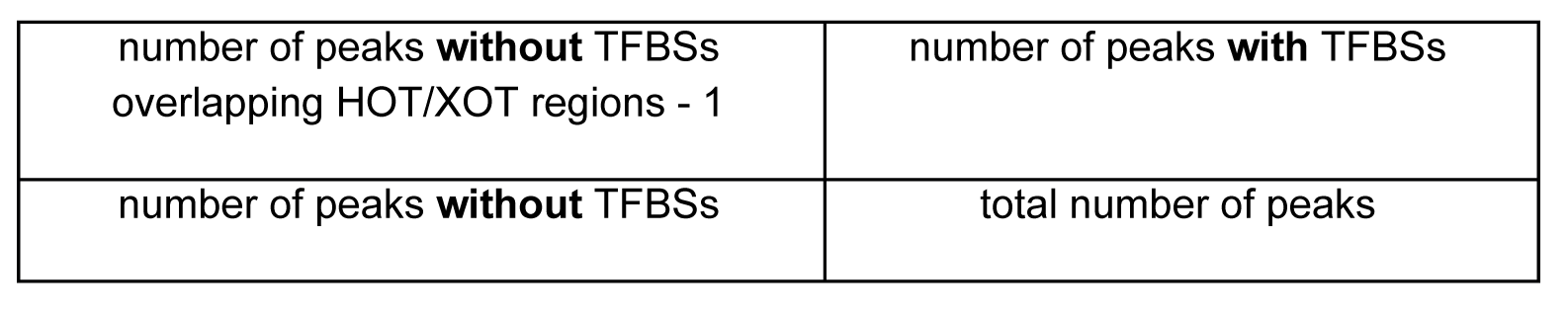

### Identification of TFs with co-localized TFBSs

For each pair of distinct TFs (TF_*A,*_, TF_B_), we extracted the closest TFBS associated with TF_B_ for each TFBS associated with TF_A_ and computed the geometric mean distance between midpoints of the paired TFBSs. With this approach, the geometric mean *m*_*AB*_ for the pair (TF_A_, TF_B_) is different from the geometric mean of the pair (TF_B_, TF_A_). With 232 TFs available in our analyses, we computed geometric means for 53,592 ordered pairs of TFs.

The colocalization of TFBSs for each TF pair was assessed using a Monte Carlo-based approach as follows. The number of TFBSs per TF ranged from 1 to 404,566, with 455 as the fifth percentile. We uniformly discretized the range [455, 414,172] to consider 50 TFBS set sizes (*S*_*i*_ for *i* in [1, 50]). We chose 414,172 as the maximum value to be able to compute a p-value for the set of 404,566 TFBSs. For each set size *S*_*i*_, we created 500 sets of TFBS by randomly selecting TFBSs from the total pool. Using these random sets, we computed null distributions for 500 Monte Carlo samples of geometric mean distances for each of the 2,601 set size combinations. Specifically, this computation led to 2,601 distributions of 500 geometric means. For the TF pair (TF_A_, TF_B_) with *N*_*A*_ and *N*_*B*_ TFBSs, respectively, we extracted the Monte Carlo sample of geometric mean distances *M* obtained from the random sets with *S*_*A*_ and *S*_*B*_ TFBSs, where *S*_*A*_ = min(*S*_*i*_) with *S*_*i*_ > *N*_*A*_ and *S*_*B*_ = min(*S*_*i*_) with *S*_*i*_ > *N*_*B*_. The empirical p-value associated with the pair (TF_A_, TF_B_) was computed as the number of times we observed a geometric mean smaller than *m*_*AB*_ from *M* over the 500 pre-computed geometric means; if no smaller geometric mean was observed, the empirical p-value is defined as < 0.002 (that is 1 / 500).

Since the expected geometric mean distance increases with a decreasing number of TFBSs, this p-value computation is conservative (under-estimated significance). The obtained p-values were corrected for multiple testing using the Benjamini-Hochberg method (52), only the TF pairs with a FDR < 5% were considered significant.

The detailed null distribution values can be downloaded and reproduced at https://hyperbrowser.uio.no/geirksa_sandbox/u/gsandve/h/null-distributions-for-manuscript-a-map-of-direct-tf-dna-interactions-in-the-human-genome. These computations are based on running the static methods “ConcatenateNullDistributionsTool.execute” and “ComputeNullDistributionForEachCombinationFromSuiteVsSuiteTool.execute” (with argument values corresponding to parameter settings annotated in the Galaxy (53) history above) in the code provided at https://hyperbrowser.uio.no/geirksa_sandbox/static/hyperbrowser/files/div/hb.zip. The source code for the comparison with null distributions is available at https://bitbucket.org/CBGR/co-binding/.

### GeneMANIA

We used the GeneMANIA software (54) to extract known protein-protein interactions from the list of TFs with significant co-localized TFBSs and plot the corresponding network.

### Prediction of *cis-*regulatory modules

The TFBSs predicted by ChIP-eat were sorted and merged using the bedtools *sort* and *merge* subcommands. The CREAM tool (55) was applied to the merged TFBSs to define *cis*-regulatory modules (CRMs) as genomic regions enriched for clusters of TFBSs.

### GWAS trait- and disease-associated single nucleotide polymorphism enrichment analysis

We assessed the enrichment for GWAS trait- and disease-associated single nucleotide variants (SNPs) at CRMs using the *traseR* R package (version 1.10.0; (56)). CRM genomic positions were lifted over to the hg19 version of the human genome to perform the analyses. The set of SNPs (as of April 30, 2018) considered by *traseR* combined data from dbGaP (57) and NHGRI (58) as described in the corresponding bioconductor package vignette (https://bioconductor.org/packages/release/bioc/vignettes/traseR/inst/doc/traseR.pdf).

### Conservation analysis

The hg38 phastCons (59) scores for multiple alignments of 99 vertebrate genomes to the human genome were retrieved as a bigWig file at http://hgdownload.cse.ucsc.edu/goldenpath/hg38/phastCons100way/hg38.phastCons100way.bw. The TFBSs predicted by ChIP-eat were sorted and merged using the bedtools *sort* and *merge* subcommands. The locations overlapping CRMs were obtained using the bedtools *intersect* subcommand. The corresponding genomic locations (for all TFBSs and TFBSs in CRMs) in BED format were decomposed into 1 bp intervals using bedops v.2.4.14 (60) with the --chop 1 option. The phastCons scores at every bp were extracted with the *ex* subcommand of the bwtool (61) using the corresponding BED and phastCons bigWig files.

### The UniBind web interface

All the TFBS predictions, corresponding ReMap ChIP-seq peaks, trained TFBS computational models, and CRMs are available through the UniBind database at http://unibind.uio.no/. The UniBind web interface was developed in Python using the model-view-controller framework Django. It uses MySQL to store TFBS metadata and Bootstrap as the frontend template engine. The source code is available at https://bitbucket.org/CBGR/unibind.

### Statistical analyses

All statistical analyses were performed in the R environment (version 3.4.4).

## RESULTS

### Predicting direct TF-DNA interactions in the human genome from ChIP-seq data

Given a set of ChIP-seq peaks and a TFBS computational model such as a PWM, one can extract the best site per peak, which corresponds to the DNA subsequence of the peak with the highest score for the model. The higher the score, the stronger the computational evidence that the site is similar to TFBSs known to be bound by the TF (34). Moreover, it has been shown that the closer the site to the peak summit, the more likely it is to represent a direct TF-DNA interaction with experimental evidence from the ChIP-seq assay (23, 25, 28). Hence, direct TF-DNA interactions captured by ChIP-seq are enriched for high scores and small distances to the peak summits (Figure 1A,B). These characteristics have previously been used to automatically predict direct TF-DNA interactions by selecting score and distance thresholds defining these enrichment zones using a heuristic approach (23). This approach used pre-defined parameter values and was specifically designed for PWMs, but is not applicable to more recent TFBS computational models such as binding energy models (BEMs) (30), transcription factor flexible models (TFFMs) (29), and DNA shape-based models (DNAshapedTFBS) (12).

**Figure 1:**
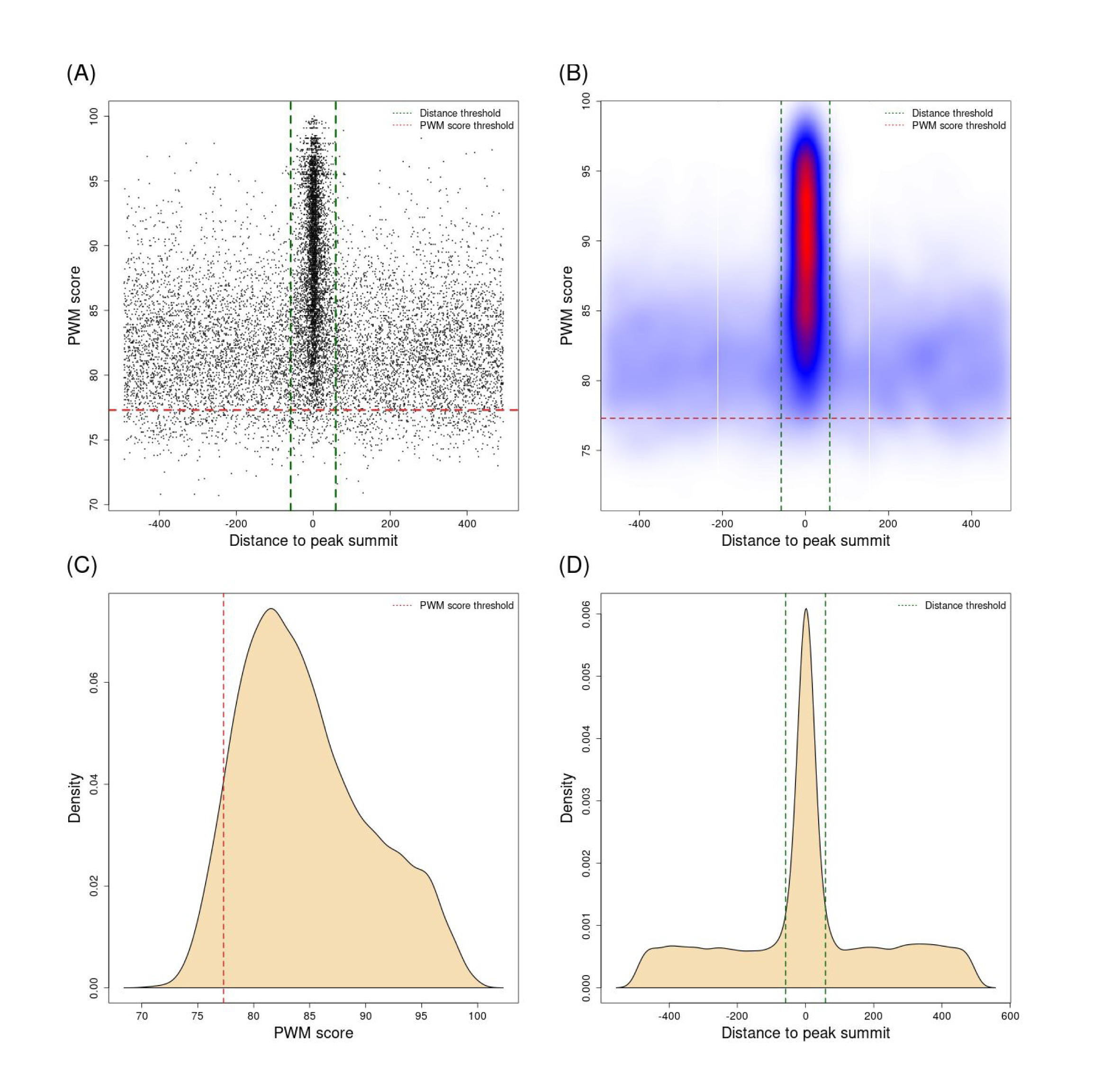
Automatic detection of the TFBSs enrichment zone. Landscape plots (23) obtained with SRF ChIP-seq peaks using the DiMO-optimized PWM MA0083.3 from JASPAR are presented as scatter (**A**) and heatmap (**B**) plots. The enrichment zone (defined within the red and green dashed line boundaries, A-B) is automatically obtained by ChIP-eat with thresholds on PWM scores (red dashed lines; **C**) and distances to peak summits (green dashed lines; **D**). The enrichment zone provides TFBSs in ChIP-seq peaks (points in A) with supporting evidence for direct TF-DNA binding from the ChIP-seq assay (close distance to peak-summits, A-B, x-axis) and the computational model (PWM score, A-B, y-axis). Distances to peak summits in A, B, and D are provided using a base pair unit.

We aimed to predict direct TF-DNA interactions (TFBSs) within ChIP-seq peaks and developed the ChIP-eat software that automatically identifies the enrichment zone for any TFBS computational model. It uses a non-parametric, entropy-based algorithm originally designed to separate background/noise from foreground/signal in image processing (40) (Supplementary Figure S2). We applied this algorithm to the distributions of site scores and distance to peak summits independently to separate direct TF-DNA interaction events from other binding subtypes and ChIP-seq artifacts (Figure 1C,D; Materials and Methods). The two thresholds define the enrichment zone, which delimits the sites that are predicted as TFBSs with both experimental and computational evidence of direct TF-DNA interactions. With this approach, we automatically adjust the enrichment zone discovery specifically for each TF ChIP-seq peak data set and for each computational model. The identified enrichment zone defines the thresholds on the TFBS computational model scores and distances to the peak summits in a data set-specific manner.

We retrieved 1,983 ChIP-seq peak data sets from ReMap (13), accounting for 232 TFs with a PFM available in the JASPAR database (21). Using DiMO-optimized PWMs, we compared the enrichment zones predicted by ChIP-eat with the ones obtained with the heuristic approach developed in (23). The enrichment zones predicted with ChIP-eat were more stringent than with the heuristic algorithm (Supplementary Figure S3A,B,D,E). The corresponding TFBSs predicted in the enrichment zones were more central to the peak summits with ChIP-eat than with the heuristic method as evaluated with CentriMo (25) (Supplementary Figure S3C,F). Moreover, ChIP-eat does not require any fixed values such as a predefined bin size (23) to predict the enrichment zones. Finally, ChIP-eat is not restricted to work with PWMs only and can be used with any TFBS computational model.

We applied ChIP-eat to the 1,983 human ChIP-seq data sets with four types of computational TFBS models: DiMO-optimized PWMs, BEMs, TFFMs, and DNAshapedTFBS. These models were optimized for each ChIP-seq data set, independently (see Materials and Methods). In the following analyses, we focused on the predictions obtained with the DiMO-optimized PWMs (see Materials and Methods). This set of direct TF-DNA interactions (TFBSs) extracted from the enrichment zones covers about 4% of the human genome, encompassing 8,304,135 distinct TFBS locations.

### Predicted direct TF-DNA interactions are likely bona fide TFBSs

#### Robustness of the enrichment zone identification

Considering the ChIP-seq data sets for the 10 most frequently ChIP’ed TFs, we observed that the thresholds on the PWM scores and distances to peak summits, defining the enrichment zones, were consistent between data sets for the same TF (Figure 2A,B). Namely, the median pairwise difference between PWM score thresholds for the same TF ranged from 1.7 to 3.7 and the median distance thresholds from 12 bp to 35 bp. As expected, the thresholds identified for distinct TFs are different (Figure 2C,D). Taken together, these results highlight that the entropy-based algorithm allows for the identification of enrichment zones specific to each TF and ChIP-seq data set, with consistent predictions between data sets for the same TF. Results were consistent with BEM, TFFM, and DNAshapedTFBS models (Supplementary Figures S4-S6).

**Figure 2:**
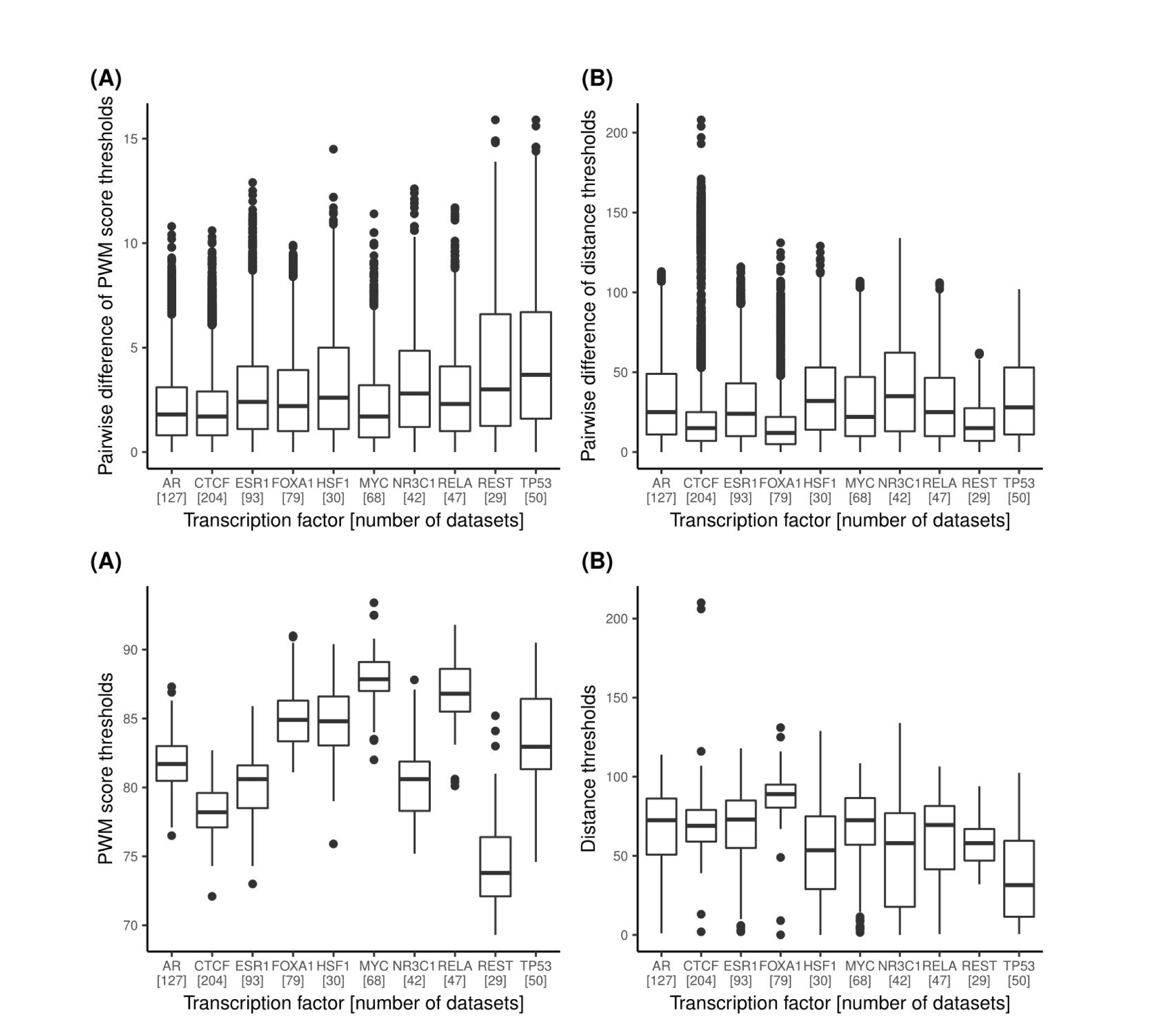
Assessment of the thresholds predicted by ChIP-eat across data sets. Boxplots of the pairwise differences for DiMO-optimized PWM score thresholds and distances to peak summits thresholds between ChIP-seq data sets for the same TF are provided in panels (**A**) and (**B**), respectively. Absolute variations of DiMO-optimized PWM score thresholds and distances to the peak summits within all data sets for the same TF are provided in panels (**C**) and (**D**), respectively. The ten TFs with the highest number of data sets were selected; the number of data sets for each TF is provided between brackets.

Further, we evaluated the robustness of the method to noise by adding 10%, 25%, and 50% of shuffled sequences to the initial set of ChIP-seq peaks for all ChIP-seq peak data sets (see Materials and Methods). The median threshold on the distances to peak summits shifted from 64 bp in the initial set of ChIP-seq peaks to 62 bp with 10% noise, 59 bp with 25% noise, and to 55 bp when adding 50% noise. The median PWM score threshold was 79 for the initial set of ChIP-seq peaks and shifted to 78.6 when adding 10% of noise, to 78.3 when adding 25% of noise, and to 78 when adding 50% of noise. A visual representation for the 10 most frequently ChIP’ed TFs is available in Supplementary Figure S7. The variability of the thresholds defining the enrichment zones when adding noise is limited, within the range of variability between ChIP-seq peak data sets for the same TF (Figure 2). Taken together, these results show that the entropy-based thresholding algorithm delimiting the enrichment zones, as implemented in ChIP-eat, is robust to random noise.

#### Validation using in vitro DNA binding affinities

To confirm *a posteriori* the high quality of our set of TFBS predictions, we assessed the TF binding affinity to DNA sequences derived experimentally from protein binding microarrays (PBM) (62). The PBM assay quantifies the binding affinity of a protein to all possible combinations of 8-mer DNA sequences. We retrieved PBM data from the UniPROBE database (45) for 40 different TFs present in our collection, corresponding to 249 ChIP-seq data sets (Supplementary Table S2). Note that the JASPAR PFMs for the ATF1, ATF3, and FOXJ2 TFs were originally derived from PBM data. For each ChIP-seq data set, we tested if the sites located in the enrichment zone presented higher binding affinity than sites outside (see Materials and Methods). The distributions of the binding affinity scores for sites within and outside the enrichment zones were compared using a Mann-Whitney signed-rank test (Figure 3A; Materials and Methods). Predicted direct TF-DNA interactions (sites within the enrichment zone) had significantly higher binding affinity than the other sites for 75% of the data sets with p-value < 0.01 and 81% with p-value < 0.05 (Figure 3B). This analysis emphasizes that the sites predicted in the defined enrichment zones are likely to correspond to direct TF-DNA interactions.

**Figure 3:**
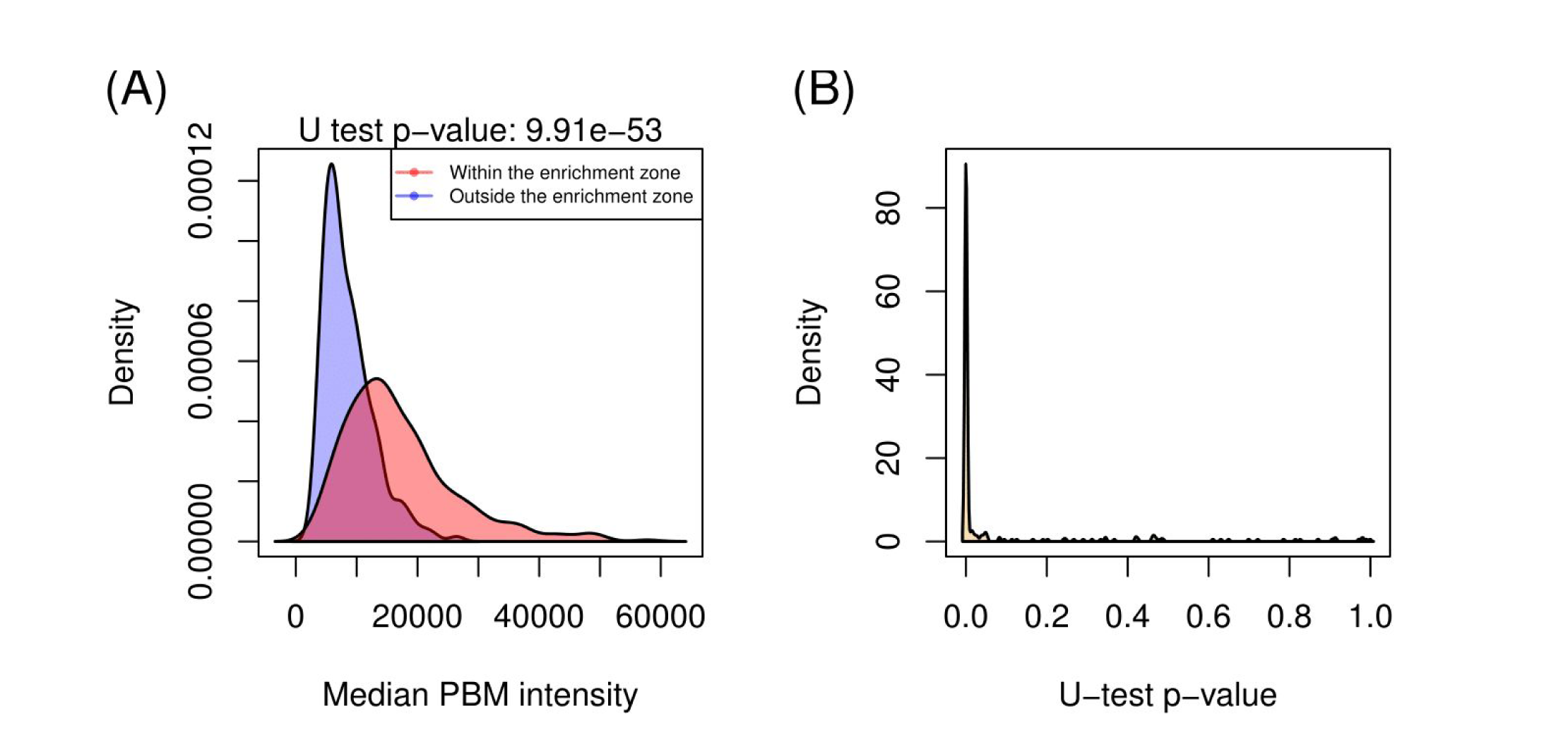
Binding affinity assessment of the predicted direct TF-DNA interactions. (**A**) Distribution of the median PBM intensity scores for the ENCSR000BMX GATA3 ChIP-seq data set between sequences at TFBSs (i.e. sites within the enrichment zone; in red) and sites outside the enrichment zone (in blue). (**B**) Distribution of Mann-Whitney U test p-values across the 249 data sets, showing distinct distributions of PBM intensity scores between sites within and outside the enrichment zones.

#### Predicted direct TF-DNA interactions are found in high confidence ChIP-seq peaks

We hypothesized that the ChIP-seq signal at ChIP-seq peaks containing a predicted direct TF-DNA interaction were more likely to be higher than at the other peaks. To test this hypothesis, we looked at (i) the quality of the peaks based on p-values assigned to the peaks by the MACS2 peak-caller and (ii) the reproducibility of calling these peaks with multiple peak-callers (MACS2, HOMER, and BCP; see Materials and Methods).

We observed that the distribution of p-values assigned by MACS2 to the peaks containing a predicted TFBS were significantly (p-value < 0.01; Mann-Whitney signed-rank test) lower than for the rest of the peaks for 1862 (96%) data sets (Figure 4). The other 77 data sets contained a reduced number of peaks (median of 837 compared to 18,968 for the complete set of ChIP-seq data sets), which can explain the lack of statistical significance. These results confirm that the predictions of direct TF-DNA interactions were found in ChIP-seq peaks of higher quality as assessed by MACS2.

**Figure 4:**
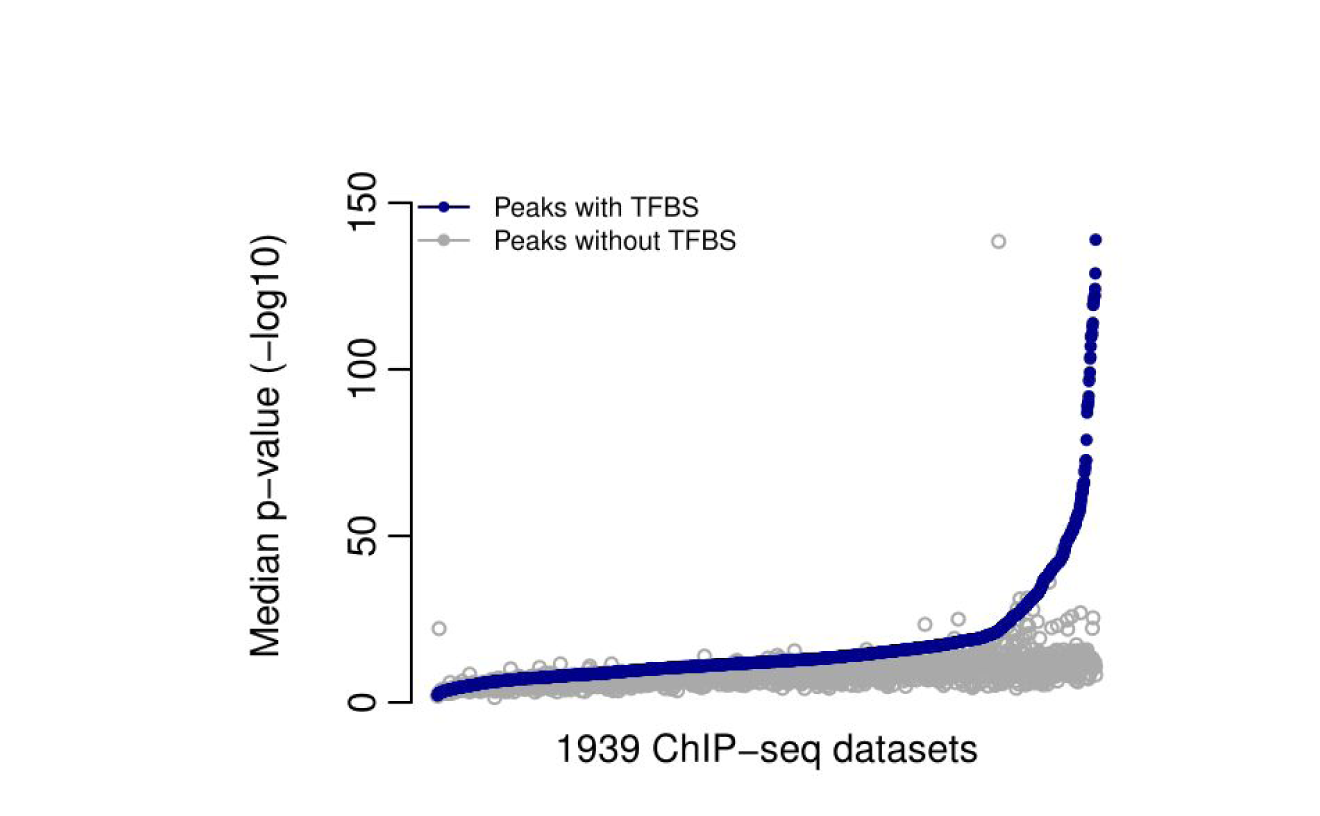
Quality assessment of the ChIP-seq peaks derived from direct TF-DNA interactions. Distribution of the median MACS2 p-values (y-axis) of the across all data sets. Values for peaks containing a predicted TFBSs are provided in blue and values for the other peaks in grey. 1,939 ChIP-seq data sets were predicted to contain direct TF-DNA interactions.

To test ChIP-seq peak-calling reproducibility, we used two other peak-callers (HOMER and BCP) on 670 ChIP-seq data sets from ENCODE. Our choice of peak-callers was motivated by their distinct statistical approaches for peak prediction. While MACS2 and HOMER are based on an empirical model supported by a Poisson distribution, BCP uses a Bayesian approach implementing infinite-state hidden Markov models. We applied ChIP-eat to the ChIP-seq peaks to predict TFBSs. For each pair of peak-callers, we assessed whether the peaks predicted to contain a direct TF-DNA interaction were more prevalent (p-value < 0.01, hypergeometric test) in the set of peaks called by both peak-callers. This was observed for 63% of the data sets for MACS2 and BCP, 70% for MACS2 and HOMER, and 66% for HOMER and BCP. The data sets without significant enrichment had a median number of peaks predicted to be derived from direct TF-DNA interactions that was ∼7 fold smaller (e.g. 3358 compared to 22,499 between MACS2 and BCP) than for the data sets with significant enrichment, and a median number of peaks without TFBS ∼2 fold larger (e.g. 40,050 compared to 21,256 between MACS2 and BCP) (Supplementary Table S3). Moreover, the median quality scores assigned by the peak-callers to the peaks from the enriched data sets were significantly (p-value < 0.01, Mann-Whitney U test) higher than for the peaks in the other data sets (Supplementary Figure S8). It suggests that the data sets enriched for reproducible peaks containing predicted direct TF-DNA interactions are of better quality than the rest of the data sets.

Taken together, these results highlight that the ChIP-seq peaks in which ChIP-eat predicts direct TF-DNA interactions are of higher quality than the other peaks. Note that the ChIP-eat tool does not consider the peak quality when predicting direct TF-DNA interactions. These observations reinforce the confidence in the predicted TFBSs by ChIP-eat.

### Predictions of direct TF-DNA interactions in ChIP-exo data

The ChIP-exo assay has been developed to provide a higher resolution than ChIP-seq to identify TFBSs *in vivo (32)*. We aimed at assessing the performance of ChIP-eat on predicting direct TF-DNA interactions using ChIP-exo data. The ChExMix tool has recently been introduced to characterize protein-DNA binding event subtypes from ChIP-exo peaks (47). ChExMix predicted different binding event subtypes for ChIP-exo data obtained for the TFs ESR1 and FOXA1, one of these subtypes corresponding to direct TF-DNA interactions (47). We applied ChIP-eat on the same ESR1 and FOXA1 ChIP-exo data sets. We compared the set of peaks identified to contain direct TF-DNA interactions predicted by ChExMix and ChIP-eat in these two data sets. We found that 93.6% (for ESR1) and 91.3% (for FOXA1) of the peaks predicted to contain TFBSs by ChIP-eat were also predicted as direct binding events by ChExMix (Supplementary Table S4). The high overlaps between the predictions from ChExMix and ChIP-eat were confirmed by Jaccard similarity indexes of 63.7% and 68.7% for ESR1 and FOXA1, respectively. The similar results obtained with the two tools suggest that ChIP-eat, designed for the more noisy and less precise ChIP-seq data, is able to capture direct binding events from ChIP-exo data.

### High-occupancy target regions are likely not derived from direct TF-DNA interactions

High-occupancy target (HOT) and extreme-occupancy target (XOT) regions are genomic regions where ChIP-seq peaks were observed for a large number of distinct ChIP’ed TFs (33, 63, 64). These regions are observed across species (64) and contain an unusually high frequency of ChIP-seq peaks (33, 63, 64). We used our set of high quality TFBS predictions to confirm that HOT/XOT regions were depleted of direct TF-DNA interactions. Indeed, we found that ChIP-seq peaks that do not contain a predicted TFBS were significantly enriched at HOT/XOT regions (odds ratio = 1.43 for HOT and 1.44 for XOT, p-value < 2.2e-16, hypergeometric test, Supplementary Table S5). This observation, combined with a previous study describing that HOT/XOT regions are likely to be derived from ChIP-seq artifacts (6), suggests that HOT/XOT regions are not derived from the direct binding of TFs.

### Predicted direct TF-DNA interactions reveal co-binding TFs and cis-regulatory modules enriched for disease- and trait-associated SNPs

TFs are known to collaborate through specific co-binding at *cis*-regulatory modules (CRMs) to achieve their function (1, 34). Hence, identifying co-binding TFs is critical to decipher transcriptional regulation of gene expression. We aimed at using our predicted direct TF-DNA interactions to reveal co-binding TFs and CRMs. We hypothesized that the distances between TFBSs of cooperating TFs are smaller than expected by chance. We tested this hypothesis for all pairs of TFs for which we predicted TFBSs (232 TFs, 53,592 pairs tested; see Materials and Methods). For each TF pair, we used a conservative Monte Carlo-based approach to compare the geometric mean of the distances between their TFBSs to the geometric mean distance expected by chance for a similar number of TFBSs randomly selected from the complete pool of TFBSs (see Materials and Methods). This approach predicted 150 pairs of TFs (accounting for 112 distinct TFs) with TFBSs closer in the genome than expected by chance (FDR < 5%; Supplementary Table S6). For 82% of the predicted TF pairs, we confirmed that the corresponding TFs physically interact using the protein-protein interaction networks from the GeneMANIA tool (54) (Supplementary Figure S9). This analysis further supports the biological relevance of the TFBSs predicted by ChIP-eat.

Next, we aimed to automatically identify CRMs, which correspond to clusters of direct TF-DNA interactions, using the clustering of genomic regions analysis method (CREAM; (55)). When considering our complete set of TFBSs, CREAM detected 61,934 CRMs in the human genome, encompassing 2,474,587 distinct TFBS locations. We found that the predicted CRMs were significantly enriched (FDR-corrected p-value = 2.9e^-150^) for disease- and trait-associated SNPs using traseR (56). Further, we observed that the TFBSs lying within the CRMs were more conserved that TFBSs predicted outside (Supplementary Figure S10). Taken together, these results indicate a potentially functional role of the CRMs identified as clusters of direct TF-DNA interactions.

### The UniBind web interface to access our collection of direct TF-DNA interactions

We catalogued the complete set of TFBS predictions from each prediction model, trained models, original ChIP-seq peaks from ReMap, and computed CRMs, and made them publicly available through UniBind at http://unibind.uio.no/. UniBind provides an interactive web interface with easy browsing, searching, and downloading for all our predictions (Figure 5). For instance, users can search for predictions for specific TFs, cell lines, and conditions.

**Figure 5:**
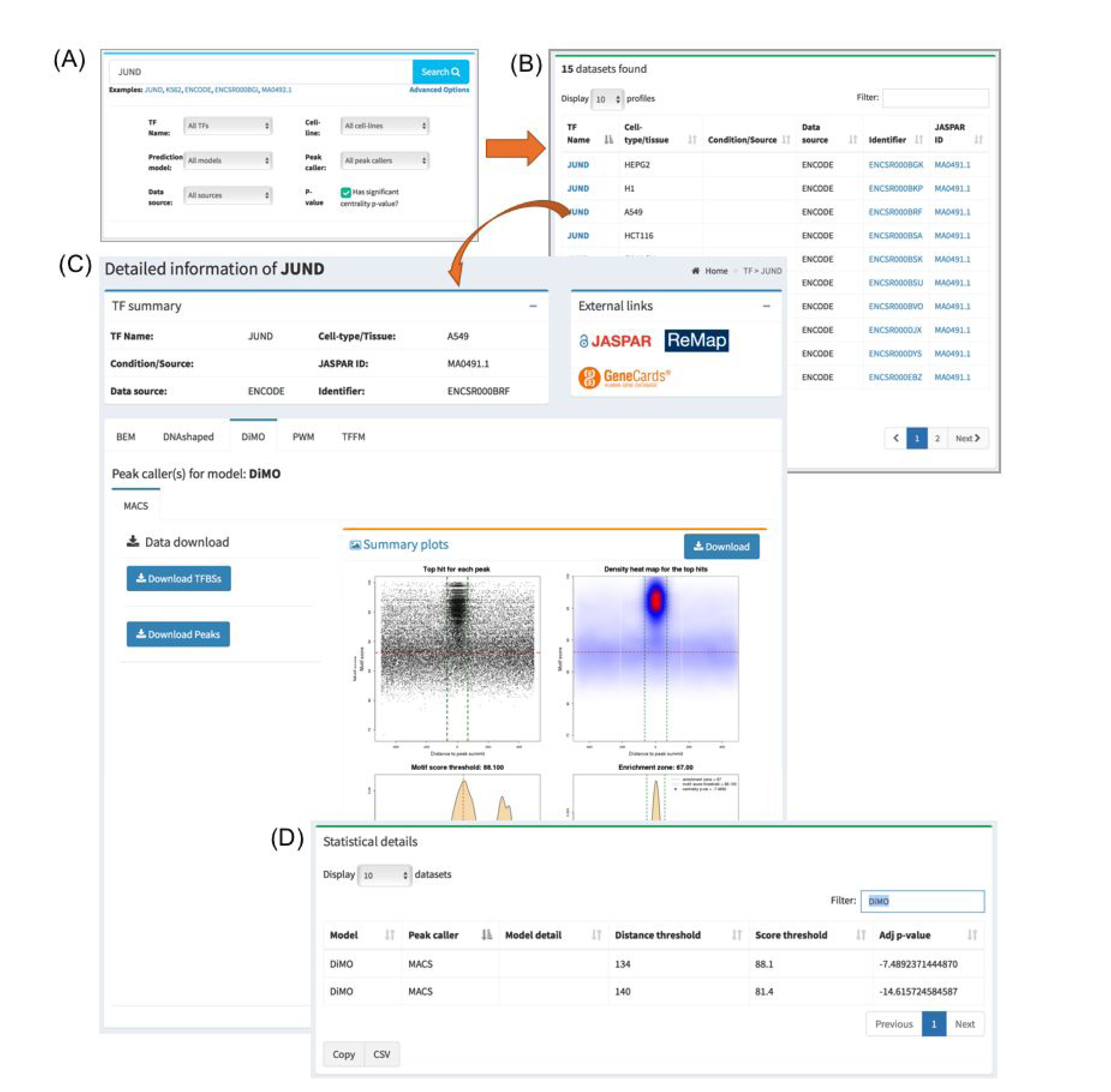
Overview of the UniBind user interface with interactive searching activity. (**A**) A quick and detailed search feature on the homepage. (**B**) A responsive table lists the searched data set(s), which can be clicked to view the details. (**C**) A detailed page shows the analysis for the JUND TF in cell-line A549, which is divided into sub-panels including the TF summary, external links, summary plots, and download options for each computational TFBS model. (**D**) Statistical details of the results.

The data can be searched by using the case insensitive search option available on the homepage. The database can be searched for each of the four TF binding models, cell/tissue type, and TF name using the ‘Advanced Options’, available on the homepage (Figure 5A). Search results are presented in a responsive and paginated table along with metadata information (Figure 5B), which can be clicked to view the detail information and download TFBSs, summary plots, and ReMap ChIP-seq peaks (Figure 5C-D). All the metadata in the responsive tables can be downloaded as CSV files. UniBind displays by default the results obtained with the DiMO-optimized PWMs but results obtained from all TFBS computational models along with the trained models are available for browsing and/or download.

## DISCUSSION

To summarize, we have uniformly processed 1,983 ChIP-seq peak data sets to predict high quality direct TF-DNA binding interactions in the human genome. The predictions were obtained using a non-parametric, entropy-based algorithm that automatically detects thresholds for TFBS computational model scores and distances to peak summits for each ChIP-seq data set. This new approach identified TFBSs supported by strong experimental and computational evidences for direct TF-DNA interactions. The accuracy of the predictions was *a posteriori* validated using the PBM *in vitro* assay, ChIP-exo data, and multiple ChIP-seq peak-calling algorithms. Our set of direct TF-DNA interactions confirmed that HOT genomic regions are likely not derived from direct binding of the TFs to the DNA. We used our TFBSs to predict TFs with proximal binding events in the human genome, which could cooperate to achieve specific functions. Further, we defined *cis*-regulatory modules, which are clusters of TFBSs, that were enriched for disease- and trait-associated SNPs from GWAS. The complete set of predictions is publicly and freely available through the UniBind web-interface (http://unibind.uio.no/), in an effort to provide the community with an unprecedented collection of high quality direct TF-DNA interaction events in the human genome.

The output of ChIP-seq assays is generally composed of direct protein-DNA interactions, indirect binding of the protein to the DNA (through a co-binding partner), nonspecific protein binding to the DNA, and noise/bias/artifacts (4, 5, 7). Here, we specifically aimed at identifying direct TF-DNA interaction events by using an entropy-based algorithm (40). This algorithm was originally developed to discriminate between foreground and background in image processing. Hence, it assumes the presence of background (or noise) in the data. As a consequence, our approach is limited by the assumption that there is background/noise in the ChIP-seq data sets analyzed. We assume that this noise represents indirect binding of TFs, nonspecific binding, or ChIP-seq experimental artifacts. Moreover, our approach considered the best site per ChIP-seq peak (defined using TFBS computational models), which represents the best candidate. We recognize that other sites with lower scores could represent direct TF-DNA interactions. These limitations denote that our approach is stringent for the prediction of direct TF-DNA interactions, favoring specificity over sensitivity. The ChIP-seq peaks that our method did not predict to contain direct TF-DNA binding events could be further analyzed to discriminate other mechanisms for protein-DNA interactions from background noise, as proposed in the ChExMix tool established for ChIP-exo data (47).

The ChIP-eat pipeline developed for this study used four TFBS computational models to predict TF-DNA binding events. These models were specifically trained for each ChIP-seq data set to improve the quality of the predictions, as the best-performing computational model varies for different TFs or TF families (9, 11, 12). As a consequence, we advocate that a ‘one-fits-all’ TFBS prediction model is not optimal and that one should compare results from multiple models. With the predictions available through UniBind, users can assess which model would perform better for each data set. Of course, it requires to use a specific metric to compare performance. As our methods aimed at identifying enrichment zones centered around ChIP-seq peak summits, we suggest to rely on a centrality measure as implemented in the CentriMo method (25). In UniBind, we provide centrality p-values computed following (25) for the predictions from each model in each ChIP-seq data set. Moreover, the ChIP-eat pipeline is generalizable and users can incorporate other TFBS computational models to predict direct TF-DNA interactions and compare them to the ones already stored in UniBind.

While studies alike focus on determining where TFs directly interact with DNA, our understanding of how these TF-DNA interactions influence expression is limited. Surely, it is critical to decipher the relationship between TF-DNA interactions and transcriptional regulation (65). It is expected that a large portion of the TFBSs identified in our study are not functional, as suggested by the futility theorem (34). Nevertheless, functional TF binding events are likely to be clustered (66–69) and associated with stronger ChIP-seq peak signals (10, 70). We expect that the direct TF-DNA interactions predicted in *cis*-regulatory modules and stored in UniBind are more likely to be enriched for functional events. Determining the specific set of functional TF-DNA interactions would require dedicated computational models and experiments.

## AVAILABILITY

Source code of the ChIP-eat software is available at https://bitbucket.org/CBGR/chip-eat_public and of UniBind at https://bitbucket.org/CBGR/unibind. The source code used for the identification of co-localized TFs is available at https://bitbucket.org/CBGR/co-binding. Users can browse and/or download the data through the UniBind web interface at http://unibind.uio.no/.

## ACKNOWLEDGEMENT

As research parasites (71), we would like to thank all the researchers who deposited their data. We thank Georgios Magklaras and his team for systems support, Andrea Cremaschi and Manuela Zucknick for statistical insights, Elisa Bjørgo and Ingrid Kjelsvik for management support, and Roza Berhanu Lemma, Jaime Castro-Mondragon, Oriol Fornes, and Phillip Richmond for comments on the manuscript draft.

